# Does reduced reactivity explain altered postural control in Parkinson’s disease? A predictive simulation study

**DOI:** 10.1101/2025.04.30.651447

**Authors:** Julian Shanbhag, Sophie Fleischmann, Iris Wechsler, Heiko Gassner, Jürgen Winkler, Bjoern M. Eskofier, Anne D. Koelewijn, Sandro Wartzack, Jörg Miehling

**Affiliations:** Engineering Design, Department of Mechanical Engineering, Friedrich-Alexander-Universität Erlangen-Nürnberg, Germany; Machine Learning and Data Analytics Lab, Department Artificial Intelligence in Biomedical Engineering (AIBE), Friedrich-Alexander-Universität Erlangen-Nürnberg, Germany; Department of Molecular Neurology, Universitätsklinikum Erlangen, Friedrich-Alexander-Universität Erlangen-Nürnberg, Germany; Digital Health and Analytics, Fraunhofer Institute for Integrated Circuits IIS, Erlangen, Germany; Chair of Autonomous Systems and Mechatronics, Department of Electrical Engineering, Friedrich-Alexander-Universität Erlangen-Nürnberg, Germany

**Keywords:** Postural control, Parkinson’s disease, neuromus-culoskeletal modelling, predictive simulation, biomechanics

## Abstract

Postural instability represents one of the cardinal symptoms of Parkinson’s disease (PD). Still, internal processes leading to this instability are not fully understood. Simulations using neuromusculoskeletal human models could help understand these internal processes leading to PD-associated postural deficits. In this paper, we investigated whether reduced reactivity amplitudes resulting from impairments due to PD can explain postural instability as well as increased muscle tone as often observed in individuals with PD. To simulate reduced reactivity, we gradually decreased previously optimized gain factors within the postural control circuitry of our model performing a quiet upright standing task. After each reduction step, the model was again optimized. Simulation results were compared to experimental data collected from 31 individuals with PD and 31 age- and sex-matched healthy control participants. Analyzing our simulation results, we showed that muscle activations increased with a model’s reduced reactivity, as well as joint angles’ ranges of motion (ROMs). However, sway parameters such as center of pressure (COP) path lengths and COP ranges did not increase as observed in our experimental data. These results suggest that a reduced reactivity does not directly lead to increased sway parameters, but could cause increased muscle tone leading to subsequent postural control alterations. To further investigate postural stability using neuromusculoskeletal models, analyzing additional internal model parameters and tasks such as perturbed upright standing requiring comparable reaction patterns could provide promising results. By enhancing such models and deepening the understanding of internal processes of postural control, these models may be used to assess and evaluate rehabilitation interventions in the future.

**Impact Statement:** Altered postural control in Parkinson’s disease can be partly associated to reduced reactivity. Additional changes in the neural circuitry need to be further investigated to fully explain the observed differences.

## I. INTRODUCTION

Upright standing is an inherently unstable body pose requiring control by the central nervous system (CNS) through specific muscle reactions, a neuromuscular process called postural control. Due to neurological disorders, such as Parkinson’s disease (PD), postural control impairment is common during the disease course. The internal processes underlying both healthy and impaired postural control in PD still remain not fully understood, with postural instability resulting from such impairments. While experimental measurements analyze PD-associated changes in postural control, they have to focus on accessible postural control outcomes and cannot provide insights into internal neural processes. Forward dynamic simulations using musculoskeletal models can help to investigate and understand these internal postural control processes by replicating these neural circuits.

Postural instability is a cardinal symptoms of PD that worsens with disease progression and increases fall risk [1]. This instability is evident in reduced stability margins and increased postural sway compared to healthy individuals [2]– [7]. Differences are observed in parameters such as increased sway range or area. Furthermore, postural reactions to external perturbations show to be smaller and slower (bradykinetic) compared to healthy persons [1]. Even individuals with PD without diagnosed balance dysfunctions can already exhibit altered sway parameters [5].

To analyze upright standing balance, forward dynamic postural control simulations can be used. Current research shows considerable variation in simulation detail, depending on the scope. Shanbhag *et al*. [8] reviewed modelling approaches that used different biomechanical models, sensor information, and control strategies. Many studies use torque-actuated single, double or triple inverted pendulum models to investigate postural control [9]–[14]. These models simplify the body’s degrees of freedom (DOFs), focusing on lower limb motion, as most postural control reactions emerge from the ankle and hip joints, eventually complemented by knee joint movements [15]. However, in order to address physiological and pathophysiological postural control aspects in more detail, musculoskeletal models with more DOFs could provide additional insights. Koelewijn *et al*. [16] and Suzuki *et al*. [17] both used musculoskeletal models incorporating proprioceptive muscle information to control upright standing. Jiang *et al*. [18] added vestibular and visual sensor information to their musculoskeletal model. Shanbhag *et al*. [19] published a detailed neuromusculoskeletal postural control model considering somatosensory, vestibular, and visual inputs, along with physiologically plausible neural delays based on sensor type and muscle position.

Up to now, only a few models have investigated aspects of postural control in the context of PD specifically, such as Boonstra *et al*. [20], Kim *et al*. [21], Dash *et al*. [22], Feller *et al*. [23], Omura *et al*. [24], and Rahmati *et al*. [25]. Of the aforementioned studies, only Omura *et al*. [24] used a musculoskeletal model to investigate postural control in individuals with PD – most models are limited to one or two DOF inverted pendulum models. However, it is essential to consider muscles in postural control simulations, as symptoms in PD, such as increased muscle tone, could independently affect other postural control aspects, such as sway behaviour. Using predictive neuromusculoskeletal simulations allows us to determine the influence of isolated parameter adaptations on postural control. Furthermore, gaining deeper insights into postural control processes in PD will enable the application of these models for assessing and evaluating rehabilitation interventions in the future.

In this paper, we investigate how reduced reactivity amplitudes affect postural control. We will use the term *reactivity* to describe the model’s reactive behaviour based on control gain factors leading to reflex amplitudes. We will answer the research question whether reducing previously optimized gain factors replicates postural control aspects associated with PD. We hypothesize that this reduction will increase sway parameters and result in higher muscle activations as a compensatory mechanism. This approach could explain reduced postural stability and higher muscle tone often observed in individuals with PD. Additionally, our investigation will provide insights into possible internal postural control processes and identify potential elements responsible for postural instability in PD. Examining the impact of reducing previously optimized control gains will reveal whether a non-ideal optimization could induce PD-associated postural control.

### II. MATERIALS AND METHODS

A previously published postural control model [19] served as a starting point to simulate upright standing postural control in the sagittal plane (section II-A). In section II-B, we describe the model adaptations to investigate whether reduced control gains replicate PD-associated motion behaviour. Forward dynamic simulations were performed using SCONE 2.3.1 [26] with Hyfydy [27] (section II-C). Experimental data from individuals with PD and healthy control (HC) participants, as described in section II-D, were used to evaluate simulations. Therefore, the analysis of the experimental data also focused on anterior-posterior (AP) movements (sagittal plane).

### A. Postural control model of upright standing

The postural control circuitry based on Shanbhag *et al*. [19] consisted of a musculoskeletal model and a neural controller comparing sensor information to a reference state resulting in specific muscle excitations, all under the influence of neural delays.

Details about the corresponding elements of this postural control model are presented in the Supplementary Materials.

### B. Simulation of pathologies

We aimed to investigate whether reducing control gains could cause PD associated motion behaviour, since reduced reactivity is often observed in individuals with PD. Reduced gain factors represent a decreased reactive response to sensory-detected deviations from a reference state. An optimization according to Shanbhag *et al*. [19] — assumed to be physiological postural control — served as a starting point. From this previously optimized simulation, we gradually reduced gain factors. We modified the model in two different ways to analyze resulting postural control. First, we reduced all gain factors equally, in the following called whole-body approach (WA). Second, we reduced only distal gain factors for lower leg muscles (gastrocnemius medialis, soleus, tibialis anterior) investigating the influence of distal elements to be more affected, in the following called distal approach (DA). Gain factors were reduced gradually (2.5 % intervals) from 100 % to 0 % for both approaches. The control model, including all remaining free parameters such as feedforward muscle excitations as well as not previously reduced gain factors (only applicable for DA), was re-optimized at each reduction step. An overview of the postural control model with its modifications to simulate a decreasing reactivity is represented in Fig. 1.

**Fig. 1.**
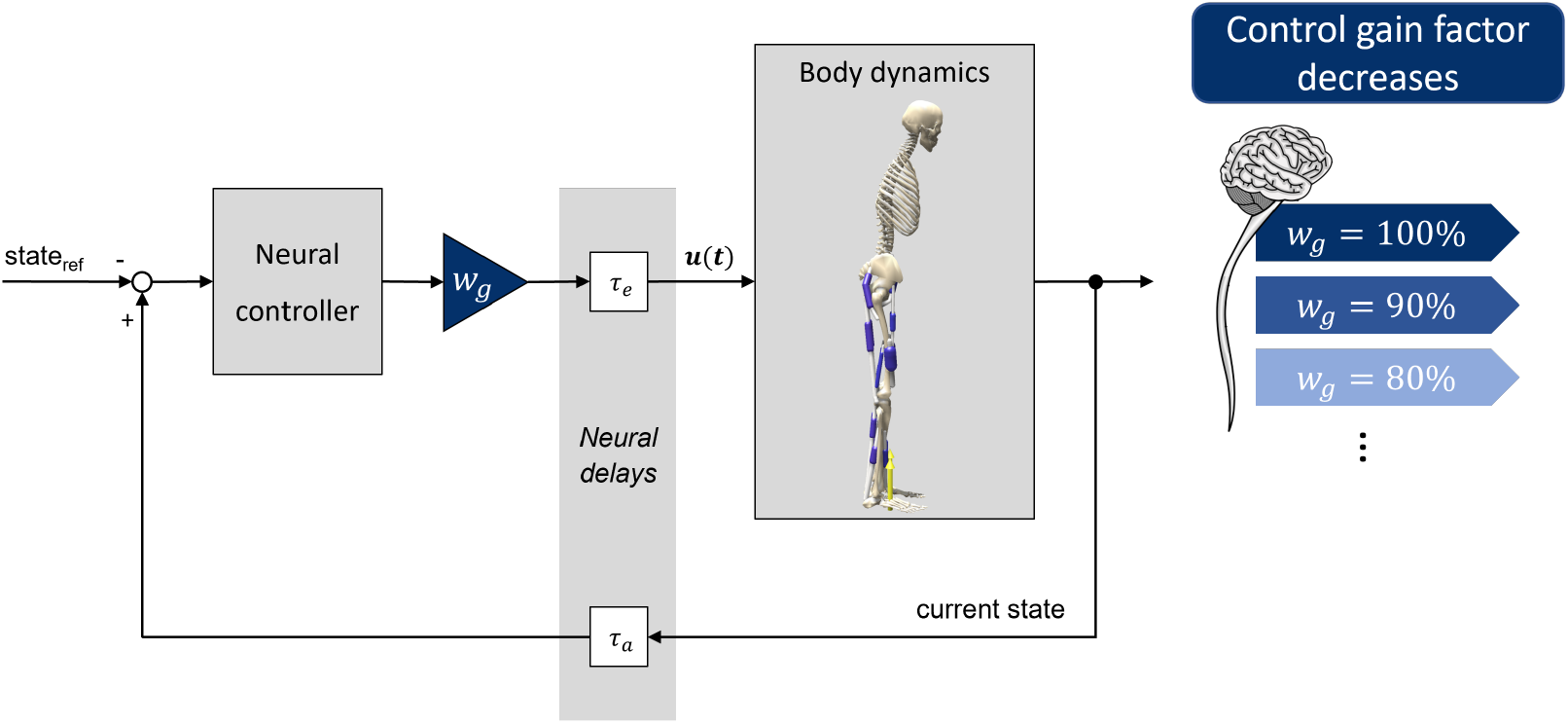
The postural control model consists of a biomechanical human model and a neural controller considering neural delays leading to muscle excitations based on the differences between current and reference states, as described in [19]. Gradually reduced gain factors, used to simulate decreasing reactivity, are incorporated through the additional factor *w*_*g*_.

### C. Simulation approach

We investigated changes in postural control by applying gradually reduced gain factors, as described in section II-B, to the following scenario: Full simulations lasted 75 seconds, evaluating the last 60 seconds due to the initial non-representative pose from default SCONE settings for standing scenarios [26]. Sampling frequency was set to 200 Hz. We tested the postural control model and its adaptations for quiet upright standing to ensure comparability to our experimental study data (section II-D).

### D. Experimental data

We conducted a study to investigate the postural control of individuals with PD and compare these study outcomes with our simulation results. 31 individuals with PD were recruited to participate in this study. Exclusion criteria were orthopedic or neurological comorbidities potentially affecting gait and balance. An age- and sex-matched HC group of 31 participants was used for comparison during the standing task.

Study population characteristics are provided in Table I. All individuals with PD performed assessments in ON medication without the presence of clinically relevant motor fluctuations or dyskinesia (one participant with active deep brain stimulation). Disease duration was defined as the time from clinical diagnosis of PD. The Hoehn & Yahr (H&Y) and UPDRS-III scores were conducted by certified clinical experts. Details regarding the statistical analyses verifying the groups’ sex- and age-matching, as well as the determining group differences in the Montreal Cognitive Assessment (MoCA) and Falls Efficacy Scale International (FES-I), are presented in the Supplementary Materials. The study was conducted in accordance with the Declaration of Helsinki and approved by the Ethics Committee of the FAU Erlangen-Nürnberg (protocol code 20-473 1-B, date of approval 09 January 2023). All participants have signed a written consent form for this study.

**TABLE I.** Characteristics of study cohort.

|  | HC group<br>(n = 31) | PD group<br>(n = 31) | p-value |
| --- | --- | --- | --- |
| Sex (male:female) | 18:13 | 25:6 | 0.0538 |
| Age (years) | 61.65 ± 13.39 | 62.65 ± 12.49 | 0.7621 |
| Disease duration (years) | – | 7.26 ± 5.18 | – |
| H&Y (score) | – | 2.13 ± 0.56 | – |
| UPDRS-III (score) | – | 20.81 ± 7.97 | – |
| MoCA (score) | 26.61 ± 2.40 | 24.42 ± 4.06 | **0.0391\*** |
| FES-I (score) | 17.53 ± 2.66 | 23.26 ± 7.58 | **0.0012\*** |

\*Significant group difference ( $p < 0.05$ ).

All participants performed an 60-second upright standing task with eyes open. Feet were placed shoulder-width apart, hands placed on the side of the hips during the whole task. The view was directed on a sign at eye level two meters in front of them. We collected the data using an optical motion capture system (VICON Vero, ten cameras, 100 Hz) and two force plates (AMTI, 1000 Hz). We used a full-body marker set consisting of 45 reflective markers according to the Plug-in Gait model [28], with additional markers on the medial knees and ankles as well as on both trochanters.

Information about the data evaluation procedure is outlined in the Supplementary Materials.

## III. RESULTS

All determined biomechanical and sway parameters for the HC and PD groups are summarized in Table II. We used the Mann-Whitney U test to compare each parameter between groups, ensuring robustness regardless of data distribution normality. By determining which parameters differed significantly between these two participant groups, comparisons of simulated results with specific experimental outcomes were feasible. Significant increases were observed in center of pressure (COP) path length, COP range, and all ranges of motion (ROMs) for pelvis, hip, knee, and ankle angles in individuals with PD compared to HC participants. No significant differences were detected for the COP position and the mean COP frequency, though a non-significant forward shift in COP position was noted in individuals with PD.

**TABLE II.** Resulting parameters from experimental data of the HC group as well as individuals with PD. All parameters focus on AP movements.

| Parameters | Control group | PD group | p-value |
| --- | --- | --- | --- |
| COP path length [mm] | 7.12 ± 2.33 | 11.58 ± 5.35 | **<0.0001\*** |
| COP range [mm] | 29.04 ± 11.80 | 38.53 ± 16.32 | **0.0163\*** |
| COP position [%] | 61.10 ± 6.17 | 62.72 ± 7.53 | 0.6495 |
| Mean COP frequency [Hz] | 0.19 ± 0.11 | 0.20 ± 0.10 | 0.5023 |
| Pelvis angle ROM [deg] | 2.36 ± 1.14 | 3.28 ± 1.73 | **0.0187\*** |
| Hip angle ROM [deg] | 3.00 ± 2.36 | 4.08 ± 2.33 | **0.0324\*** |
| Knee angle ROM [deg] | 1.70 ± 1.52 | 3.14 ± 2.38 | **0.0013\*** |
| Ankle angle ROM [deg] | 1.27 ± 0.75 | 2.18 ± 1.71 | **0.0092\*** |

\*Significant group difference ( $p < 0.05$ )

We further divided the PD group into patients with and without postural instability based on their H&Y score to get an impression about the parameters’ progression direction depending on disease severity. The parameters are visualized in Fig. 2. Due to the small number of highly affected individuals with PD (PD-PI: four with H&Y 3, one with H&Y 4), we did not further analyze these data statistically according to these subgroups. However, parameters tended to increased in the more severely affected patient subgroup.

**Fig. 2.**
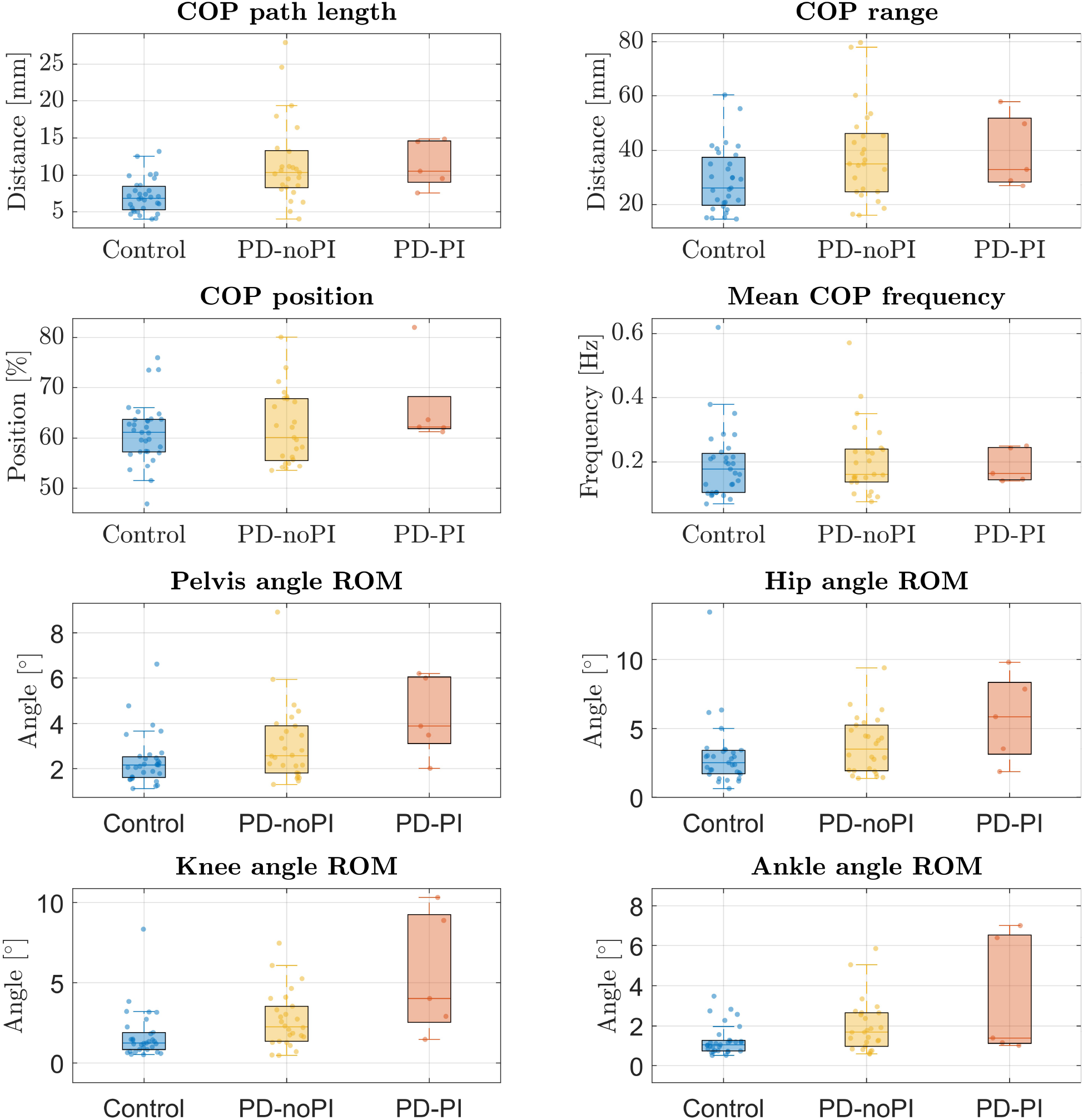
COP parameters and joint angles’ ROMs during quiet upright standing of a control group and individuals with PD, separated into patients without (PD-noPI) and with postural instability (PD-PI). Feet were placed shoulder-width apart during this standing task. All parameters focus on AP movements.

We conducted simulations with gradually reduced gain factors. The WA model was able to fulfill the upright standing task until 67.5 %, and the DA model until 57.5 % of optimal gain factors. For smaller gain factors, the model was not able to keep standing for the 75-second simulation. In the following, only successful tasks were further analyzed.

Fig. 3 represents resulting parameters for the different simulations, with specific values in Tables I and II in the Supplementary Materials. Correlations between gain factor reductions and each parameter were determined using Spearman’s correlation coefficient. Both WA and DA simulations showed heterogeneous results when reducing gain factors. The COP path lengths decreased in both WA and DA. The COP range remained unchanged for both approaches. Relative COP positions showed a forward shift with smaller gains in both, but significantly for WA only. The mean COP frequency did not change significantly for the WA, but decreased for the DA with smaller gain factors. For the WA, pelvis, hip, and ankle ROMs were increased with decreased gain factors; knee ROMs showed little effects. For the DA, only the ankle’s ROM changed remarkably depending on gain factors; however, none of the joint angle ROMs changes correlated significantly with the adapting gain factors.

**Fig. 3.**
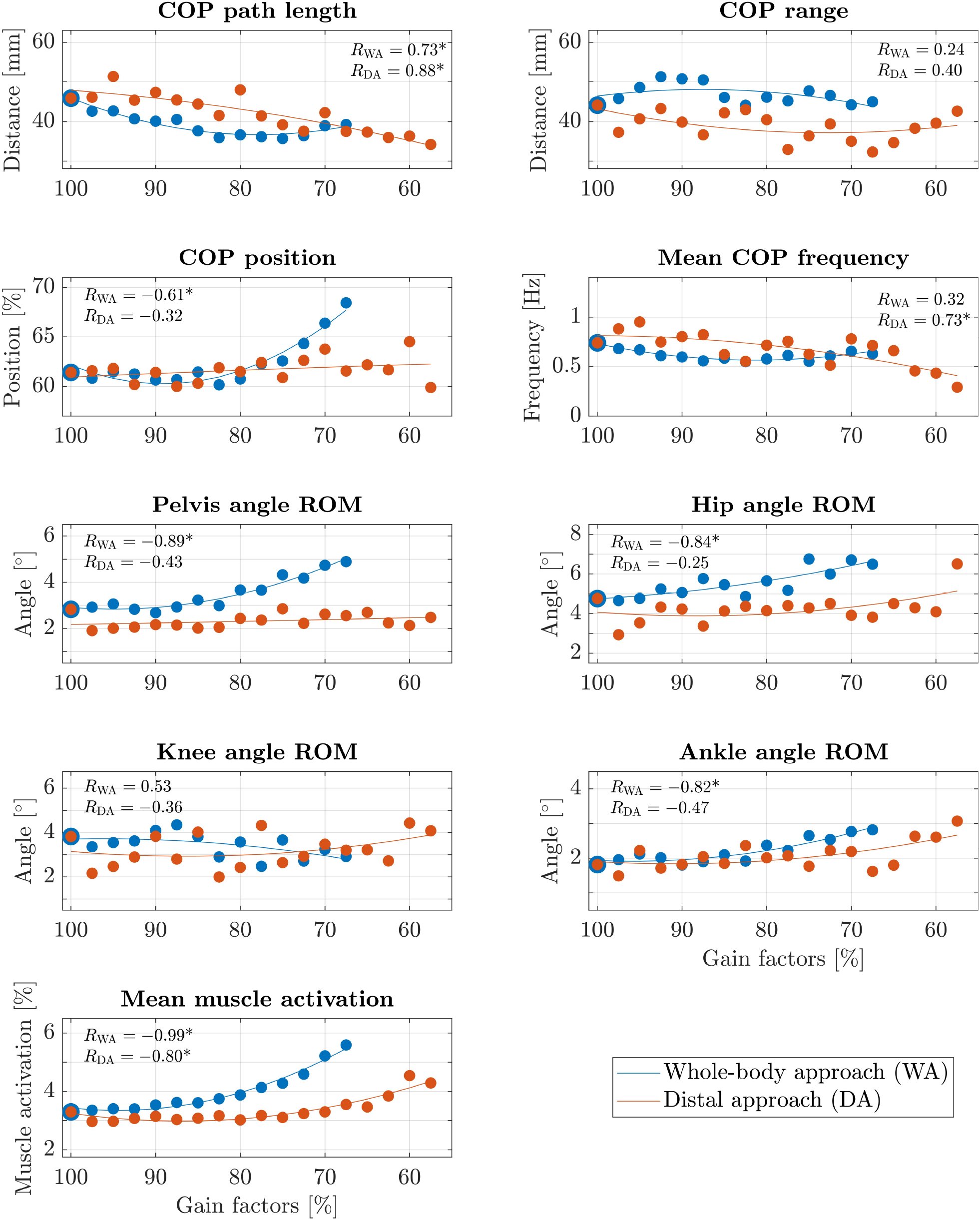
COP parameters, joint angles’ ROMs and mean muscle activations of quiet upright standing simulations depending on different gain factor reductions visualized with additional regression curves. All parameters focus on AP movements.

Both approaches (WA and DA) resulted in consistently increasing mean muscle activations with reduced gain factors.

## IV. DISCUSSION

### A. Comparison of experimental data and simulation results

We aimed to answer the research question of whether reducing gain factors in postural control simulations replicates motion aspects that can be associated with PD. We hypothesized that reducing gain factors would lead to increased sway parameters and higher muscle activations.

For both simulation approaches (WA and DA), the model exhibited significantly higher muscle activations with decreased gain factors. This finding aligns with reports about increased muscle tone in individuals with PD [7], [29], [30]. We could not directly compare this aspect to our experimental data since we did not conduct maximum voluntary contraction measurements. However, this increased muscle activity can be understood as a compensatory mechanism for the reduced reactivity to maintain balance during upright standing. These higher muscle activations also represent increased co-contractions and higher joint stiffness, which could explain the decreased sway parameters.

Simulation results showed diverse parameter trends compared to our experimental data on postural stability. Generally, parameters based on joint angles showed comparable trends, parameters based on the COP tended to vary more between experimental and simulation data. For individuals with PD compared to the HC group, and for gradually reduced gain factors in our WA simulations, we found increasing ROMs for the pelvis, hip, and ankle joints. The DA did not show significant trends for the joint angles’ ROMs. For the WA, the COP position showed a significant forward shift with decreasing gain factors, aligning with the trend observed in our experimental data, even if not statistically significant. This effect could already be seen in other experimental studies [5], [31]. This tendency was also recognized for the DA but was not significant. Aligning with our experimental data, the COP frequency showed no significant changes in WA simulations. The DA represented significantly decreasing frequencies with reducing gain factors. However, we observed higher-frequency COP movements in our overall simulation results. This observation suggests that our model may still include aspects that lead to higher frequencies in the COP. Although we observed significantly increased COP path lengths in our experimental data of individuals with PD, simulation results of both, WA and DA, showed significantly decreased COP path lengths with reduced gain factors. Our experimental data showed significantly higher COP ranges of individuals with PD; however, simulation results did not correlate with varied gain factors. Therefore, reduced reactivity cannot explain the changes in COP path length and range seen in our experimental data. However, it has to be mentioned that there are also findings from different studies describing smaller COP ranges for individuals with PD, e.g. from Horak *et al*. [32]. Reasons for these differences compared to our study outcomes could be a smaller but higher affected patient group (7 patients, average H&Y 3.4 *±* 0.2) as well as a different movement task that could significantly influence outcome parameters. Therefore, study cohorts and movement tasks should be carefully identified before comparing different study results.

Overall, the WA resulted in parameter changes more similar to experimental data of individuals with PD, suggesting that all gain factors could be affected similarly in PD. As postural control reactions are centrally processed in the body’s CNS, and degenerative processes due to PD also occur within the CNS, it is reasonable to assume that the gain factors are not influenced by the position of the respective muscle, but are rather uniformly affected in the same incorrect manner.

Differences not explained by reduced gain factors, representing the model’s decreased reactivity, could result from different elements within the neural circuit for postural control. One aspect also associated with PD is increased noise during motor tasks. Feller *et al*. [23] reported a decreased signal-to-noise ratio in processing sensory signals in individuals with PD. This additional noise might explain the higher COP path lengths and ranges observed in the PD group, which our current simulations have not confirmed yet due to uniform noise levels for all investigated scenarios. Additionally, it could be questioned, whether individuals with PD actually use the same optimization objectives, with factors like fear of falling potentially playing an important role.

### B. Findings gained from experimental data

Analyzing our experimental data of individuals with PD and an age- and sex-matched control group, we found significant differences in COP path lengths, COP ranges, as well as all joint angles’ ROMs. This aligns with many findings from literature [3], [4], [33], [34]. We did not see significant group differences in sway frequencies, based on COP data. However, other studies could observe differences in sway frequencies. Kamieniarz *et al*. [35] and Rocchi *et al*. [36] found higher mean frequencies of individuals with PD compared to a control group, whereas Kwon *et al*. [34] detected lower mean frequencies of the PD group compared to healthy age-matched participants. This discrepancy suggests that this aspect is not unambiguous and might depend on the patient group, especially the number of participants with different disease stages or symptom severity. In this respect, our patient population was quite homogeneous and did not represent many patients suffering from severe postural instability represented by an H&Y score of 3 or higher. This could explain why no significant differences in sway frequencies could be identified in our study. The same applies to the COP position. Błaszczyk *et al*. [5] and Bloem *et al*. [31] could identify a forward-shifted COP for individuals with PD. We did not see significant differences in our experimental data. However, especially in Fig. 2, a tendency towards a forward shift of the COP is prevalent.

### C. Limitations

We reduced the control gains compared to previously optimized values to simulate the effects of inadequate gain scaling. Since PD involves complex alterations within the CNS and motor control, it is plausible that changes in motion behaviour are also influenced by additional parameters within the model. In our simulations, we reduced control gains equally for every control element. Currently, sensory information type is not considered, which could be influenced differently in PD. By considering this aspect in a next step, differences that can be observed in short and long latency reflexes between individuals with PD and healthy persons could be taken into account [37], [38]. The current model only represented movements in the sagittal plane and was controlled symmetrically. However, important aspects of postural instability in PD are also observed in the body’s frontal and coronal plane [39] and show to be partly asymmetrical as well [20], [30]. We used a generic, not a person-specific scaled musculoskeletal human model to conduct our simulations. We would expect simulation results to be even more similar to experimental data using person-specific scaled models representing the study’s participants. Additionally, it is important to consider that we assumed the initial model to represent physiologically plausible postural control. Nevertheless, a model representing human-like movements does not necessarily prove internal neural control processes to exactly align with the model.

Our study consisted of 31 individuals with PD with an average H&Y of 2.13 *±* 0.56, but only five had a H&Y of 3 or 4, limiting our group with diagnosable postural instability. To observe even more distinct differences between individuals with PD and HC participants, we would recommend including more participants suffering from postural instability. We could not compare amplitudes of simulated muscle activations with experimental data, as participants did not perform maximum voluntary contraction measurements before conducting EMG data. Understanding absolute muscle activations and their changes based on disease stages would be beneficial for comparison with our simulations. Lastly, we could only measure quiet upright standing; exploring postural control in additional scenarios would be extremely insightful and could further deepen our understanding of the adapted model’s gain factors.

## V. CONCLUSION

In this paper, we investigated how reduced reactivity affects postural control using forward dynamic simulations. We found that some parameter changes associated with PD were observed in our simulation results. Especially muscle tone showed to be increased, compensating for reduced reactivity. However, other aspects, like increased COP path lengths and COP ranges, could not be observed in our simulations. This suggests that reduced reactivity alone is not the only factor causing postural instability in PD. Other elements of the postural control circuitry need to be investigated next to explain remaining differences between individuals with PD and healthy persons reflecting postural instability. By deepening our understanding of internal postural control processes, these models and predictive simulations may be used in the future for assessing and evaluating rehabilitation interventions. Once capable of reproducing common differences in motion behaviour between healthy persons and individuals with PD, models can provide tailored intervention recommendations based on a person’s previous measurements and further person-specific simulations.

## Supplementary Materials

The Supplementary Materials file includes additional information about the postural control model, the experimental data, data evaluation, and detailed tables presenting the biomechanical parameters of the simulation results.

## References

[1] J.-H. Park, Y.-J. Kang, and F. B. Horak, “What Is Wrong with Balance in Parkinson’s Disease?” Journal of Movement Disorders, vol. 8, no. 3, pp. 109–114, 2015, ISSN: 2005-940X, 2093-4939. DOI: 10.14802/jmd.15018.

[2] F. Doná, C. Aquino, J. Gazzola, et al., “Changes in postural control in patients with Parkinson’s disease: A posturographic study,” Physiotherapy, vol. 102, no. 3, pp. 272–279, 2016, ISSN: 00319406. DOI: 10.1016/j.physio.2015.08.009.

[3] S. D. Kim, N. E. Allen, C. G. Canning, and V. S. C. Fung, “Postural Instability in Patients with Parkinson’s Disease: Epidemiology, Pathophysiology and Management,” CNS Drugs, vol. 27, no. 2, pp. 97–112, 2013, ISSN: 1172-7047, 1179-1934. DOI: 10.1007/s40263-012-0012-3.

[4] T. Yamamoto, Y. Suzuki, K. Nomura, et al., “A Classification of Postural Sway Patterns During Upright Stance in Healthy Adults and Patients with Parkinson’s Disease,” Journal of Advanced Computational Intelligence and Intelligent Informatics, vol. 15, no. 8, pp. 997–1010, 2011, ISSN: 1883-8014, 1343-0130. DOI: 10.20965/jaciii.2011.p0997.

[5] J. W. Błaszczyk, R. Orawiec, D. Duda-Kłodowska, and G. Opala, “Assessment of postural instability in patients with Parkinson’s disease,” Experimental Brain Research, vol. 183, no. 1, pp. 107–114, 2007, ISSN: 0014-4819, 1432-1106. DOI: 10.1007/s00221-007-1024-y.

[6] J. Raymakers, M. Samson, and H. Verhaar, “The assessment of body sway and the choice of the stability parameter(s),” Gait & Posture, vol. 21, no. 1, pp. 48–58, 2005, ISSN: 09666362. DOI: 10.1016/j.gaitpost.2003.11.006.

[7] J. V. Jacobs, D. M. Dimitrova, J. G. Nutt, and F. B. Horak, “Can stooped posture explain multidirectional postural instability in patients with Parkinson’s disease?” Experimental Brain Research, vol. 166, no. 1, pp. 78–88, 2005, ISSN: 0014-4819, 1432-1106. DOI: 10.1007/s00221-005-2346-2.

[8] J. Shanbhag, A. Wolf, I. Wechsler, et al., “Methods for integrating postural control into biomechanical human simulations: A systematic review,” Journal of neuroengineering and rehabilitation, vol. 20, no. 1, p. 111, 2023. DOI: 10.1186/s12984-023-01235-3.

[9] Y. Li, W. S. Levine, and G. E. Loeb, “A two-joint human posture control model with realistic neural delays,” IEEE transactions on neural systems and rehabilitation engineering, vol. 20, no. 5, pp. 738–48, 2012. DOI: 10.1109/TNSRE.2012.2199333.

[10] R. J. Peterka, “Sensorimotor integration in human postural control,” Journal of neurophysiology, vol. 88, no. 3, pp. 1097–118, 2002, Num Pages: 22, ISSN: 0022-3077. DOI: 10.1152/jn.2002.88.3.1097.

[11] H. van der Kooij, R. Jacobs, B. Koopman, and F. van der Helm, “An adaptive model of sensory integration in a dynamic environment applied to human stance control,” Biological cybernetics, vol. 84, no. 2, pp. 103–15, 2001, ISSN: 0340-1200. DOI: 10.1007/s004220000196.

[12] T. A. Boonstra, A. C. Schouten, and H. van der Kooij, “Identification of the contribution of the ankle and hip joints to multi-segmental balance control,” Journal of neuroengineering and rehabilitation, vol. 10, p. 23, 2013, Num Pages: 18. DOI: 10.1186/1743-0003-10-23.

[13] C. Tian and J. He, “Simulation study of human posture control under external perturbation,” Proceedings of the 36th IEEE Conference on Decision and Control, pp. 2529–2534, 1997, Num Pages: 6. DOI: 10.1109/CDC.1997.657699.

[14] M. Günther and H. Wagner, “Dynamics of quiet human stance: Computer simulations of a triple inverted pendulum model,” Computer methods in biomechanics and biomedical engineering, vol. 19, no. 8, pp. 819–34, 2016, Num Pages: 16. DOI: 10.1080/10255842.2015.1067306.

[15] T. Mergner, “A neurological view on reactive human stance control,” Annual Reviews in Control, vol. 34, no. 2, pp. 177–198, 2010, Num Pages: 22, ISSN: 13675788. DOI: 10.1016/j.arcontrol.2010.08.001.

[16] A. D. Koelewijn and A. J. Ijspeert, “Exploring the Contribution of Proprioceptive Reflexes to Balance Control in Perturbed Standing,” Frontiers in bioengineering and biotechnology, vol. 8, p. 866, 2020, ISSN: 2296-4185. DOI: 10.3389/fbioe.2020.00866.

[17] Y. Suzuki and H. Geyer, “A Neuro-Musculo-Skeletal Model of Human Standing Combining Muscle-Reflex Control and Virtual Model Control,” 2018 40th Annual International Conference of the IEEE Engineering in Medicine and Biology Society (EMBC), vol. 2018, pp. 5590–5593, 2018. DOI: 10.1109/EMBC.2018.8513543.

[18] P. Jiang, R. Chiba, K. Takakusaki, and J. Ota, “A postural control model incorporating multisensory inputs for maintaining a musculoskeletal model in a stance posture,” Advanced Robotics, vol. 31, no. 1-2, pp. 55–67, 2017, ISSN: 0169-1864. DOI: 10.1080/01691864.2016.1266095.

[19] J. Shanbhag, S. Fleischmann, I. Wechsler, et al., “A sensorimotor enhanced neuromusculoskeletal model for simulating postural control of upright standing,” Frontiers in Neuroscience, vol. 18, p. 1 393 749, 2024, ISSN: 1662-453X. DOI: 10.3389/fnins.2024.1393749.

[20] T. A. Boonstra, A. C. Schouten, J. P. P. van Vugt, B. R. Bloem, and H. van der Kooij, “Parkinson’s disease patients compensate for balance control asymmetry,” Journal of neurophysiology, vol. 112, no. 12, pp. 3227– 39, 2014, Num Pages: 13, ISSN: 0022-3077. DOI: 10.1152/jn.00813.2013.

[21] S. Kim, F. B. Horak, P. Carlson-Kuhta, and S. Park, “Postural feedback scaling deficits in Parkinson’s disease,” Journal of neurophysiology, vol. 102, no. 5, pp. 2910–20, 2009, Num Pages: 11, ISSN: 0022-3077. DOI: 10.1152/jn.00206.2009.

[22] R. Dash, V. V. Shah, and H. J. Palanthandalam-Madapusi, “Explaining Parkinsonian postural sway variabilities using intermittent control theory,” Journal of biomechanics, vol. 105, p. 109 791, 2020, Num Pages: 8, ISSN: 0021-9290. DOI: 10.1016/j.jbiomech.2020.109791.

[23] K. J. Feller, R. J. Peterka, and F. B. Horak, “Sensory Reweighting for Postural Control in Parkinson’s Disease,” Frontiers in human neuroscience, vol. 13, p. 126, 2019, Num Pages: 17, ISSN: 1662-5161. DOI: 10.3389/fnhum.2019.00126.

[24] Y. Omura, H. Togo, K. Kaminishi, et al., “Analysis of abnormal posture in patients with Parkinson’s disease using a computational model considering muscle tones,” Frontiers in computational neuroscience, vol. 17, p. 1 218 707, 2023, Num Pages: 13, ISSN: 1662-5188. DOI: 10.3389/fncom.2023.1218707.

[25] Z. Rahmati, S. Behzadipour, A. C. Schouten, and G. Taghizadeh, “A Postural Control Model to Assess the Improvement of Balance Rehabilitation in Parkinson’s Disease,” in 2018 7th IEEE International Conference on Biomedical Robotics and Biomechatronics (Biorob), Num Pages: 6, 2018, pp. 1019–1024. DOI: 10.1109/BIOROB.2018.8487884.

[26] T. Geijtenbeek, “SCONE: Open Source Software for Predictive Simulation of Biological Motion,” Journal of Open Source Software, vol. 4, no. 38, p. 1421, 2019. DOI: 10.21105/joss.01421.

[27] T. Geijtenbeek, The Hyfydy Simulation Software, 2021.

[28] Vicon Motion Systems Limited UK, Vicon Plug-in Gait Reference Guide, 2021.

[29] A. Burleigh, F. Horak, J. Nutt, and J. Frank, “Levodopa Reduces Muscle Tone and Lower Extremity Tremor in Parkinson’s Disease,” Canadian Journal of Neurological Sciences/Journal Canadien des Sciences Neurologiques, vol. 22, no. 4, pp. 280–285, 1995, ISSN: 0317-1671, 2057-0155. DOI: 10.1017/S0317167100039470.

[30] W. Wright, V. Gurfinkel, J. Nutt, F. Horak, and P. Cordo, “Axial hypertonicity in Parkinson’s disease: Direct measurements of trunk and hip torque,” Experimental Neurology, vol. 208, no. 1, pp. 38–46, 2007, ISSN: 00144886. DOI: 10.1016/j.expneurol.2007.07.002.

[31] B. R. Bloem, D. J. Beckley, M. P. Remler, R. A. Roos, and J. Van Dijk, “Postural reflexes in Parkinson’s disease during ‘resist’ and ‘yield’ tasks,” Journal of the Neurological Sciences, vol. 129, no. 2, pp. 109–119, 1995, ISSN: 0022510X. DOI: 10.1016/0022-510X(94)00253-K.

[32] F. B. Horak, D. Dimitrova, and J. G. Nutt, “Direction-specific postural instability in subjects with Parkinson’s disease,” en, Experimental Neurology, vol. 193, no. 2, pp. 504–521, 2005, ISSN: 00144886. DOI: 10.1016/j.expneurol.2004.12.008.

[33] J.W. Błaszczyk and R. Orawiec, “Assessment of postural control in patients with Parkinson’s disease: Sway ratio analysis,” Human Movement Science, vol. 30, no. 2, pp. 396–404, 2011, ISSN: 01679457. DOI: 10.1016/j.humov.2010.07.017.

[34] D.-Y. Kwon, Y.-H. Choi, Y.-R. Kwon, G.-M. Eom, and J.-W. Kim, “Comparison of Static Postural Balance in Patients with SWEDDS and Parkinson’s Disease,” Journal of Mechanics in Medicine and Biology, vol. 20, no. 09, p. 2 040 013, 2020, Num Pages: 12, ISSN: 0219-5194. DOI: 10.1142/S0219519420400138.

[35] A. Kamieniarz, J. Michalska, W. Marszałek, et al., “Detection of postural control in early Parkinson’s disease: Clinical testing vs. modulation of center of pressure,” en, PLOS ONE, vol. 16, no. 1, J. L. McKay, Ed., e0245353, 2021, ISSN: 1932-6203. DOI: 10.1371/journal.pone.0245353.

[36] L. Rocchi, L. Chiari, A. Cappello, and F. B. Horak, “Identification of distinct characteristics of postural sway in Parkinson’s disease: A feature selection procedure based on principal component analysis,” Neuroscience Letters, vol. 394, no. 2, pp. 140–145, 2006, ISSN: 03043940. DOI: 10.1016/j.neulet.2005.10.020.

[37] P. Mazzoni, B. Shabbott, and J. C. Cortes, “Motor Control Abnormalities in Parkinson’s Disease,” Cold Spring Harbor Perspectives in Medicine, vol. 2, no. 6, a009282–a009282, 2012, ISSN: 2157-1422. DOI: 10.1101/cshperspect.a009282.

[38] B. R. Bloem, D. J. Beckley, and J. G. Van Dijk, “Are automatic postural responses in patients with Parkinson’s disease abnormal due to their stooped posture?” Experimental Brain Research, vol. 124, no. 4, pp. 481– 488, 1999, ISSN: 0014-4819, 1432-1106. DOI: 10.1007/s002210050644.

[39] M. Mancini, P. Carlson-Kuhta, C. Zampieri, J. G. Nutt, L. Chiari, and F. B. Horak, “Postural sway as a marker of progression in Parkinson’s disease: A pilot longitudinal study,” Gait & Posture, vol. 36, no. 3, pp. 471–476, 2012, ISSN: 09666362. DOI: 10.1016/j.gaitpost.2012.04.010.

