## Supplementary Materials for "Does reduced reactivity explain altered postural control in Parkinson’s disease? A predictive simulation study"

#### I. Postural control model of upright standing

*Model elements*

The musculoskeletal human model was based on Delp *et al.* [1] with updates from Rajagopal *et al.* [2] and is distributed with SCONE. The planar sagittal plane model consisted of six segments (trunk-pelvis, upper legs, lower legs, feet) with three degree of freedom (DOF) per leg and three DOF in the pelvis with respect to the ground. Motion behaviour was assumed to be symmetrical, therefore left and right muscles are controlled identically. This resulted in a total of six DOF for the model. For each leg, nine Hill-type muscles [3] were considered: Gluteus maximus, iliopsoas, hamstrings, biceps femoris short head, rectus femoris, vastus intermedius, gastrocnemius medialis, soleus, and tibialis anterior. We used a generic model as we focused on general rather than person-specific movement aspects.

The neural controller calculated muscle excitations for each muscle based on somatosensory, vestibular, visual sensor information feedback, as well as feedforward elements of constant muscle excitations. Translational and rotational movements of the head in space as well as lengths, lengthening velocities and forces from the muscles were considered as sensor information. All sensor information was compared to a reference state and the resulting differences weighted by specific gain factors. These gain factors reacting on differences between current and reference state were optimized according to section I. The resulting weighted control feedback served as muscle excitations that were applied to the model's muscles, leading to muscle activations determined via activation and contraction dynamics of the Hill-type muscle model [3].

Detailed information on this postural control model can be found in Shanbhag *et al.* [4].

*Enhanced internal perturbations*

The previous model [4] showed lower ranges of motion (ROMs) compared to experimental data, since only internal perturbations in the form of random Gaussian noise were considered. Therefore, we added low-frequent disturbances to the model representing breathing (equation 1) and heartbeat (equation 2) which both also influence postural control [5].

$$F_{\text{breathing}}(t) = A_{\text{breathing}} \cdot \sin(2\pi f_{\text{breathing}} \cdot t) \quad (1)$$

We assumed an amplitude $A_{\text{breathing}}$ of 1 N and a breathing frequency $f_{\text{breathing}}$ of 0.25 Hz (corresponding to 15 breaths per minute), resulting in a perturbation force $F_{\text{breathing}}(t)$ for continuous breathing.

$$F_{\text{heartbeat}}(t) = \begin{cases} A_{\text{heartbeat}} \cdot \sin\left(2\pi \frac{1}{\tau_{\text{pulse}}} \cdot t_{\text{rel}}\right), & \text{if } t_{\text{rel}} < \tau_{\text{pulse}} \\ 0, & \text{else} \end{cases} \quad (2)$$

$$\text{with } t_{\text{rel}} = t - n \cdot T;\ T = \frac{1}{f_{\text{heartbeat}}}$$

$F_{\text{heartbeat}}(t)$ described the perturbation force caused by the heartbeat. We assumed an amplitude $A_{\text{heartbeat}}$ of 0.1 N and a pulse duration $\tau_{\text{pulse}}$ of 100 ms representing the QRS complex of the heartbeat [6]. $t_{\text{rel}}$ represented the relative time within one period $T$, $n$ the number of periods passed, the heartbeat frequency $f_{\text{heartbeat}}$ was set to 1.1 Hz (corresponding to 66 beats per minute).

The sum of these two perturbation forces $F_{\text{perturbation}}$ was applied to the model's torso acting in the anterior-posterior (AP) direction according to equation 3.

$$F_{\text{perturbation}} = F_{\text{breathing}} + F_{\text{heartbeat}} \quad (3)$$

The individual perturbation forces as well as the resulting sum are represented in Fig. 1.

Additionally, internal noise was applied to the model's muscle excitations in the form of random Gaussian noise, assuming that there is a deviation between the ideal reaction determined by the control system and the actually executed muscle excitations.

$$u' = u + k_{noise} \cdot R \quad (4)$$

$$k_{noise} = 0.01 + 0.1 \cdot u \quad (5)$$

The resulting muscle excitation $u'$ was calculated from the ideal signal $u$, a signal-dependent noise amplitude $k_{noise}$, and a randomly generated Gaussian-distributed number $R$. The noise amplitude consisted of a base element and an element proportional to the signal's amplitude $u$. The specific amplitudes and ratios of base and proportional noise were set manually.

*Optimization of control parameters*

We applied the pre-implemented covariance matrix adaptation evolution strategy (CMA-ES) algorithm [7] in SCONE to optimize free parameters of the postural control model. Optimization criteria were applied similarly to Shanbhag *et al.* [4]. Control model parameters from the previous simulation — the scenario assumed to represent physiological postural control — served as initial guesses for the following simulations of whole-body approach (WA) and distal approach (DA).

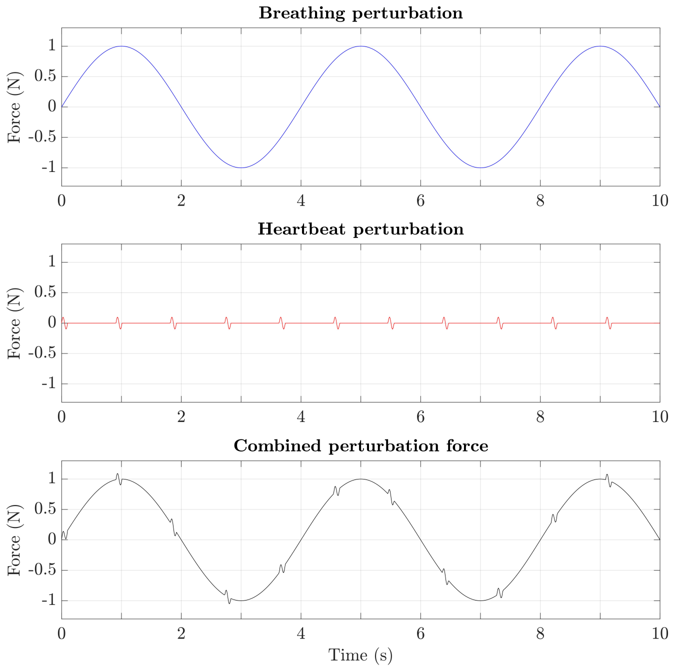


Fig. 1. Perturbation forces consisting of a breathing and a heartbeat perturbation. The resulting sum was applied to the model's torso in AP direction.

For each optimization scenario, six parallel optimizations were carried out with different random seeds. The optimizations were prioritized based on their predicted fitness values [8]. As soon as the reduction of the cost function's result, determined by an average of the last 500 generations, was smaller than 1e-5 compared to the average of the previous optimization step, the optimization ended leading to one optimized simulation result for this specific scenario.

### II. Experimental data

To verify that both participant groups were sex- and age-matched, the following statistical analyses were conducted:
A Chi-square test was used to evaluate differences in sex distribution between the groups ($p = 0.0538$; no significant difference between groups). To compare age, a Shapiro-Wilk test confirmed normal distribution($p > 0.05$), and a Levene's test indicated homogeneity of variance ($p > 0.05$). Consequently, an independent t-test was conducted, indicating that there was no significant difference in age between the two groups ($p = 0.7621$).
Furthermore, we evaluated the differences in Montreal Cognitive Assessment (MoCA) and Falls Efficacy Scale International (FES-I) between the two groups. Since both scores were not normally distributed (Shapiro-Wilk test, $p < 0.05$), a Mann-Whitney U test was conducted for each score. The results showed that both MoCA and FES-I were significantly different between the healthy control (HC) and Parkinson's disease (PD) groups ($p = 0.0391$ and $p = 0.0012$, respectively). The PD group exhibited mild cognitive impairments, as indicated by a MoCA score of $24.42 \pm 4.06$ (with scores $< 26$ suggesting cognitive impairments) [9]. Additionally, this group reported a high level of concern about falling, reflected by a FES-I score of $23.26 \pm 7.58$ (with scores $> 23$ indicating a high concern about falling) [10]. In contrast, the HC group showed no cognitive impairments and only low concerns about falling, with a MoCA score of $26.61 \pm 2.40$ and a FES-I score of $17.53 \pm 2.66$.

*Preparation for data evaluation:* Recorded marker and force plate data were pre-filtered with a third-order Butterworth filter with a cutoff frequency of 10 Hz. We used OpenSim 4.4 [11], MATLAB R2022a [12], and a three-dimensional model based on Rajagopal *et al.* [2] to process the data further. The model consisted of eight segments and 16 DOF. We scaled the model to match the anthropometrics of the study participants using an automated scaling tool [13]. Afterwards, we conducted an inverse kinematics analysis via OpenSim. Left and right joint angles were averaged as movements were assumed to be symmetrical.

In a first step, we analyzed our experimental data grouped into HC participants and individuals with PD. In a second step, we further divided the patient group, based on their Hoehn & Yahr (H&Y) score, into patients without postural instability (H&Y 1 and 2, in total 26 participants), later called PD-noPI and patients with postural instability (H&Y 3 and 4, in total 5 participants), later called PD-PI. For evaluating and comparing simulation and experimental results, we calculated biomechanical and sway parameters describing the postural stability during the upright standing task. To gain center of pressure (COP) results, data from both force plates were fused to one resulting COP for each time step by determining a weighted sum. Only AP movement components were analyzed since simulations were conducted with our sagittal plane model and did not represent movements in the frontal or coronal plane.

### III. Data evaluation

For both, experimental data and simulation results, we derived biomechanical and sway parameters to describe postural stability.

We determined the COP path length $l_{COP,AP}$ describing the total travelled distance of the COP in AP direction for the whole standing task, normalized to 1 second:

$$l_{COP,AP} = \frac{1}{T} \sum_{i=1}^{n-1} |(COP_{AP}(i+1) - COP_{AP}(i))| \tag{6}$$

$COP_{AP}$ represents the COP position in the AP direction, $T$ the total simulation time, $n$ the number of sampling points.

We calculated the mean COP position, relative to the support area described by the area underneath and between the feet:

$$p_{COP,AP} = \frac{\|COP_{AP} - M_{HEEL,AP}\|}{\|M_{TOE,AP} - M_{HEEL,AP}\|} \tag{7}$$

$M_{HEEL,AP}$ represents the heel marker position, $M_{TOE,AP}$ the toe marker position, both in the AP direction.

The mean frequency of the COP was calculated as follows:

$$\bar{f}_{COP,AP} = \frac{\sum_{i=2}^{N/2} (f_i \cdot P_i)}{\sum_{i=2}^{N/2} P_i} \tag{8}$$

$P_i$ represents the power amplitude of the power signal $P$ at the specific frequency $f_i$. The power signal was previously generated by transferring the COP signal to the frequency domain using a fast Fourier transform (FFT) in Matlab.

Additionally, COP ranges as well as ROMs of the several joint angles were calculated.

### IV. Simulation results

Tables I and II show biomechanical parameters of the simulation results that are described in section III of the main manuscript.

TABLE I. Resulting parameters from the base simulation as well as gain-reduced simulations of whole-body approach (WA). All parameters focus on AP movements. Correlations are calculated between gradually reduced gain factors and each biomechanical parameter.

| Parameters | Base simulation | Gain factors WA | | | | | | | | | | | | | | | | | Correlations |
|---|---|---|---|---|---|---|---|---|---|---|---|---|---|---|---|---|---|---|---|
| | | 97.5% | 95% | 92.5% | 90% | 87.5% | 85% | 82.5% | 80% | 77.5% | 75% | 72.5% | 70% | 67.5% | 65% | 62.5% | 60% | 57.5% | |
| COP path length [mm] | 45.90 | 42.62 | 42.69 | 40.70 | 40.10 | 40.53 | 37.67 | 35.95 | 36.65 | 36.18 | 35.72 | 36.43 | 38.96 | 39.25 | N/A | N/A | N/A | N/A | **0.73*** |
| COP range [mm] | 44.16 | 45.82 | 48.61 | 51.31 | 50.79 | 50.52 | 46.09 | 44.13 | 46.15 | 45.24 | 47.74 | 46.58 | 44.24 | 45.00 | N/A | N/A | N/A | N/A | 0.24 |
| COP position [%] | 43.77 | 43.17 | 43.82 | 43.60 | 43.00 | 43.03 | 43.81 | 42.51 | 43.10 | 44.64 | 44.93 | 46.68 | 48.74 | 50.79 | N/A | N/A | N/A | N/A | **-0.61*** |
| Mean COP frequency [Hz] | 0.74 | 0.68 | 0.67 | 0.61 | 0.60 | 0.56 | 0.58 | 0.55 | 0.58 | 0.61 | 0.55 | 0.61 | 0.65 | 0.63 | N/A | N/A | N/A | N/A | 0.32 |
| Pelvis angle ROM [deg] | 2.83 | 2.92 | 3.06 | 2.83 | 2.68 | 2.93 | 3.23 | 2.99 | 3.67 | 3.66 | 4.32 | 4.18 | 4.73 | 4.89 | N/A | N/A | N/A | N/A | **-0.89*** |
| Hip angle ROM [deg] | 4.76 | 4.66 | 4.76 | 5.24 | 5.06 | 5.77 | 5.46 | 4.86 | 5.65 | 5.17 | 6.75 | 5.99 | 6.71 | 6.50 | N/A | N/A | N/A | N/A | **-0.84*** |
| Knee angle ROM [deg] | 3.82 | 3.36 | 3.54 | 3.62 | 4.09 | 4.34 | 3.81 | 2.90 | 3.57 | 2.47 | 3.66 | 2.71 | 3.24 | 2.92 | N/A | N/A | N/A | N/A | 0.53 |
| Ankle angle ROM [deg] | 1.81 | 1.95 | 2.12 | 2.01 | 1.80 | 1.90 | 2.10 | 1.92 | 2.37 | 2.22 | 2.65 | 2.54 | 2.77 | 2.82 | N/A | N/A | N/A | N/A | **-0.82*** |
| Mean muscle activation [%] | 3.30 | 3.35 | 3.41 | 3.40 | 3.54 | 3.62 | 3.61 | 3.75 | 3.88 | 4.13 | 4.28 | 4.59 | 5.22 | 5.59 | N/A | N/A | N/A | N/A | **-0.99*** |

*Significant correlation ($p<0.05$)

TABLE II. Resulting parameters from the base simulation as well as gain-reduced simulations of distal approach (DA). All parameters focus on AP movements. Correlations are calculated between gradually reduced gain factors and each biomechanical parameter.

| Parameters | Base simulation | Gain factors DA | | | | | | | | | | | | | | | | | Correlations |
|---|---|---|---|---|---|---|---|---|---|---|---|---|---|---|---|---|---|---|---|
| | | 97.5% | 95% | 92.5% | 90% | 87.5% | 85% | 82.5% | 80% | 77.5% | 75% | 72.5% | 70% | 67.5% | 65% | 62.5% | 60% | 57.5% | |
| COP path length [mm] | 45.90 | 46.14 | 51.38 | 45.40 | 47.39 | 45.46 | 44.42 | 41.54 | 48.02 | 41.44 | 39.20 | 37.58 | 42.25 | 37.52 | 37.34 | 35.98 | 36.34 | 34.22 | **0.88*** |
| COP range [mm] | 44.16 | 37.28 | 40.69 | 43.29 | 39.87 | 36.66 | 42.21 | 43.02 | 40.47 | 32.95 | 36.40 | 39.40 | 35.03 | 32.32 | 34.70 | 38.31 | 39.61 | 42.62 | 0.40 |
| COP position [%] | 43.77 | 43.93 | 44.16 | 42.55 | 43.77 | 42.35 | 42.66 | 44.23 | 43.84 | 44.76 | 43.25 | 44.98 | 46.12 | 43.91 | 44.54 | 44.03 | 46.87 | 42.24 | -0.32 |
| Mean COP frequency [Hz] | 0.74 | 0.88 | 0.95 | 0.75 | 0.80 | 0.82 | 0.62 | 0.55 | 0.72 | 0.76 | 0.62 | 0.51 | 0.78 | 0.71 | 0.66 | 0.46 | 0.43 | 0.29 | **0.73*** |
| Pelvis angle ROM [deg] | 2.83 | 1.91 | 2.01 | 2.06 | 2.17 | 2.15 | 2.01 | 2.05 | 2.44 | 2.37 | 2.85 | 2.22 | 2.62 | 2.56 | 2.70 | 2.24 | 2.13 | 2.48 | -0.43 |
| Hip angle ROM [deg] | 4.76 | 2.93 | 3.53 | 4.33 | 4.23 | 3.37 | 4.13 | 4.36 | 4.15 | 4.40 | 4.29 | 4.51 | 3.91 | 3.82 | 4.50 | 4.30 | 4.09 | 6.51 | -0.25 |
| Knee angle ROM [deg] | 3.82 | 2.16 | 2.47 | 2.89 | 3.83 | 2.80 | 4.02 | 1.99 | 2.42 | 4.32 | 2.64 | 2.93 | 3.47 | 3.20 | 3.22 | 2.72 | 4.42 | 4.08 | -0.36 |
| Ankle angle ROM [deg] | 1.81 | 1.49 | 2.22 | 1.71 | 1.82 | 2.04 | 1.85 | 2.36 | 2.01 | 2.07 | 1.77 | 2.22 | 2.19 | 1.62 | 1.80 | 2.63 | 2.61 | 3.07 | -0.47 |
| Mean muscle activation [%] | 3.30 | 2.97 | 2.98 | 3.08 | 3.15 | 3.04 | 3.08 | 3.17 | 3.02 | 3.18 | 3.11 | 3.24 | 3.30 | 3.55 | 3.47 | 3.84 | 4.54 | 4.29 | **-0.80*** |

*Significant correlation ($p<0.05$)

### References

[1] S. L. Delp, J. P. Loan, M. G. Hoy, F. E. Zajac, E. L. Topp, and J. M. Rosen, "An interactive graphics-based model of the lower extremity to study orthopaedic surgical procedures," *IEEE transactions on bio-medical engineering*, vol. 37, no. 8, pp. 757–67, 1990, ISSN: 0018-9294. DOI: 10.1109/10.102791.

[2] A. Rajagopal, C. L. Dembia, M. S. DeMers, D. D. Delp, J. L. Hicks, and S. L. Delp, "Full-Body Musculoskeletal Model for Muscle-Driven Simulation of Human Gait," *IEEE Transactions on Biomedical Engineering*, vol. 63, no. 10, pp. 2068–2079, 2016, ISSN: 0018-9294, 1558-2531. DOI: 10.1109/TBME.2016.2586891.

[3] M. Millard, T. Uchida, A. Seth, and S. L. Delp, "Flexing computational muscle: Modeling and simulation of musculotendon dynamics," *Journal of biomechanical engineering*, vol. 135, no. 2, p. 021 005, 2013, ISSN: 0148-0731. DOI: 10.1115/1.4023390.

[4] J. Shanbhag, S. Fleischmann, I. Wechsler, *et al.*, "A sensorimotor enhanced neuromusculoskeletal model for simulating postural control of upright standing," *Frontiers in Neuroscience*, vol. 18, p. 1 393 749, 2024, ISSN: 1662-453X. DOI: 10.3389/fnins.2024.1393749.

[5] P. A. Forbes, A. Chen, and J.-S. Blouin, "Sensorimotor control of standing balance," *Handbook of clinical neurology*, vol. 159, pp. 61–83, 2018, ISSN: 0072-9752. DOI: 10.1016/B978-0-444-63916-5.00004-5.

[6] W. F. Boron and E. L. Boulpaep, *Medical Physiology E-Book : Medical Physiology E-Book*. Chantilly, UNITED STATES: Elsevier, 2016, ISBN: 978-1-4557-3328-6.

[7] C. Igel, N. Hansen, and S. Roth, "Covariance matrix adaptation for multi-objective optimization," *Evolutionary computation*, vol. 15, no. 1, pp. 1–28, 2007, ISSN: 1063-6560. DOI: 10.1162/evco.2007.15.1.1.

[8] T. Geijtenbeek, "SCONE: Open Source Software for Predictive Simulation of Biological Motion," *Journal of Open Source Software*, vol. 4, no. 38, p. 1421, 2019. DOI: 10.21105/joss.01421.

[9] Z. S. Nasreddine, N. A. Phillips, V. Bédirian, *et al.*, "The Montreal Cognitive Assessment, MoCA: A Brief Screening Tool For Mild Cognitive Impairment," en, *Journal of the American Geriatrics Society*, vol. 53, no. 4, pp. 695–699, 2005, ISSN: 0002-8614, 1532-5415. DOI: 10.1111/j.1532-5415.2005.53221.x.

[10] K. Delbaere, J. C. T. Close, A. S. Mikolaizak, P. S. Sachdev, H. Brodaty, and S. R. Lord, "The Falls Efficacy Scale International (FES-I). A comprehensive longitudinal validation study," en, *Age and Ageing*, vol. 39, no. 2, pp. 210–216, 2010, ISSN: 0002-0729, 1468-2834. DOI: 10.1093/ageing/afp225.

[11] A. Seth, J. L. Hicks, T. K. Uchida, *et al.*, "OpenSim: Simulating musculoskeletal dynamics and neuromuscular control to study human and animal movement," *PLoS computational biology*, vol. 14, no. 7, p. 1 006 223, 2018. DOI: 10.1371/journal.pcbi.1006223.

[12] The MathWorks Inc, *MATLAB (R2022a)*, Natick, Massachusetts, United States, 2022.

[13] A. Di Pietro, A. Bersani, C. Curreli, and F. Di Puccio, "AST: An OpenSim-based tool for the automatic scaling of generic musculoskeletal models," *Computers in Biology and Medicine*, vol. 175, p. 108 524, 2024, ISSN: 00104825. DOI: 10.1016/j.compbiomed.2024.108524.
